# A pilot study of the effects of *Mycoplasma ovipneumoniae* exposure on domestic lamb growth and performance

**DOI:** 10.1101/459628

**Authors:** Thomas E. Besser, Jessica Levy, Melissa Ackerman, Danielle Nelson, Kezia Manlove, Kathleen A. Potter, Jan Busboom, Margaret Benson

## Abstract

*Mycoplasma ovipneumoniae* is a globally distributed pathogen that has been associated with pneumonia in both domestic and wild Caprinae. It is closely related to *M. hyopneumoniae*, a respiratory pathogen of swine that is associated with decreased growth rates of pigs as well as clinical respiratory disease. In order to assess the effects of *M. ovipneumoniae* on lamb performance, we generated a cohort of lambs free of *M. ovipneumoniae* by segregation of test negative ewes after lambing, then compared the growth and carcass quality traits of *M. ovipneumoniae-free* and -colonized lambs from weaning to harvest. Some signs of respiratory disease were observed during the feeding trial in both lamb groups, but the *M. ovipneumoniae-exposed* group included more affected lambs and higher average disease scores. At harvest, lungs of lambs in both groups showed few grossly visible lesions, although the *M. ovipneumoniae-exposed* group did exhibit increased microscopic lung lesions (*P*<0.05). In addition, *M. ovipneumoniae* exposed lambs produced lower average daily gains (*P*<0.05), and lower yield grade carcasses (*P*<0.05) compared to those of non-exposed lambs. The results demonstrated the feasibility of test and segregation for elimination of *M. ovipneumoniae* from groups of sheep and suggested that this pathogen may impair lamb growth and productivity even in the absence of overt respiratory disease.

## Introduction

The bacterium *Mycoplasma ovipneumoniae* was originally isolated and described from a Australian sheep flock experiencing a high incidence of pneumonia in lambs [1]. Lesions associated with the disease included proliferative interstitial pneumonia with septal and bronchiolar epithelial cell hyperplasia and proliferation, with additional infiltration in some cases by lymphocytes or neutrophils. This disease has since been observed to have a global distribution, a variable but generally low severity, and has been variously termed enzootic pneumonia, chronic non-progressive pneumonia, or interstitial pneumonia [2]. In severe cases, enzootic pneumonia may involve co-infection with multiple genetic strain types of *M. ovipneumoniae* and also other bacterial respiratory pathogens, including prominently *Mannheimia haemolytica* [3, 4]. Enzootic pneumonia has been associated with adverse effects on lamb growth and productivity [5-9].

*M. ovipneumoniae* is a member of the neurolyticum/hyopneumoniae subgroup of mycoplasmas, which also includes *M. hyopneumoniae* of swine, *M. dispar* of cattle, and *M. conjunctivae* of sheep and goats [10, 11]. *M. hyopneumoniae* is recognized as an important pathogen contributing to respiratory disease and unthriftiness in pigs [12, 13]. The combined impacts of *M. hyopneumoniae* on respiratory disease and impaired growth have led to the recent development and adoption of methods to eliminate this pathogen from large swine herds through the use of breeding management combined with antibiotic therapy [14, 15]. If the effects of *M. ovipneumoniae* on lamb growth and productivity are similar to those of *M. hyopneumoniae* in swine, similar efforts at flock-level elimination of this pathogen may be merited.

*M. ovipneumoniae* is widespread in US sheep operations. The USDA National Animal Health Monitoring Service included diagnostics for *M. ovipneumoniae* in its Sheep 2011 survey of 450 US domestic sheep operations [16]. Evidence of *M. ovipneumoniae* colonization was detected in approximately 90% of operations. *M. ovipneumoniae* has also recently been identified as the primary pathogen initiating severe epizootics of pneumonia in bighorn sheep [17, 18]. Further complicating the picture, *M. ovipneumoniae* may also be detected in the lungs of lambs in the absence of overt clinical signs or grossly visible lesions of enzootic pneumonia [2, 19]. These sub-clinical infections may also play a role in impaired lamb growth and productivity that would not be detected in studies that rely solely on grossly visible lung lesions as a marker for infection.

While previous studies of enzootic pneumonia have generally compared groups of lambs that differed in the extent of grossly visible lung lesions, our approach was designed to evaluate the effects of *M. ovipneumoniae* infection regardless of the extent of visible pneumonia lesions. Therefore, the objective of this study was to determine the feasibility of producing a cohort of lambs unexposed to *M. ovipneumoniae* on an infected premises in order to evaluate the effects of *M. ovipneumoniae* infection or colonization of domestic lambs from weaning to harvest.

## Materials and Methods

### Ethics statement

This study was carried out in accordance with the recommendations in the Guide for the Care and Use of Laboratory Animals of the National Institutes of Health and in conformance with United States Department of Agriculture animal research guidelines, under protocol #04482 approved by the Washington State University Institutional Animal Care and Use Committee (WSU IACUC). Parts of this study were conducted at the University of Idaho Sheep Center under that same Washington State University IACUC protocol under a memorandum of understanding with the University of Idaho Laboratory Animal Research office.

### Animals and husbandry

The study was conducted using lambs from a cross-bred production flock of approximately 100 ewes located at the University of Idaho Sheep Center. Carrier and non-carrier ewes were identified by longitudinal testing using nasal swab samples tested by realtime polymerase chain reactions (PCR). After lambing, lambs were supplemented with selenium and vitamin E (0.5 ml Bo-Se, Merck Animal Health, Madison NJ) and carrier and non-carrier ewes with their lambs were separated and housed in non-adjacent pens to prevent nose-to-nose contact. During this period, the pre-weaned lambs had access to the feed available to the ewes, which consisted of a 50:50 mix of rolled corn and whole barley (1 kg/ewe/day) plus long-stemmed dairy-quality alfalfa hay (2 kg/ewe/day). At approximately 65 days of age the lambs were weaned, weighed, and vaccinated with Clostridial CDT bacterin/toxoid. A second dose of CDT vaccine was administered 30 d later during the feeding trial.

### Microbiology

*M. ovipneumoniae* was detected in nasal swab and lung samples using realtime PCR. Swabs (BBL CultureSwabEZ, Fisher Scientific, Waltham, MA) were serially inserted into both nares of each sampled animal, replaced in the swab sheath, and maintained at 4C for up to 3 hrs before storage at -20C until DNA extraction. At harvest, separate swabs from each principal bronchus (left, right, tracheal) were handled and stored similarly. Swabs were eluted by vigorous agitation in 400 ul of phosphate-buffered saline, centrifuged (20,000 x g, 20 min), and the supernatant was discarded. DNA was extracted from eluate pellets using Qiagen kits (Dneasy, Qiagen, Valencia CA). Realtime PCR testing was performed with a modification of a previously published method [20]. Modifications included 1) use of a new primer (226Fnew 5’-GGGGTGCGCAACATTAGTTAGTTGGTAG-3’) to replace the previously described forward primer, modified master mix to include bovine serum albumin (final concentration, 400 ng/ml), and 3) modified thermocycling conditions including Stage 1: 50C hold for 2 minutes followed by 95C hold for 10 minutes; optics off; Stage 2: 40 repeat cycles of denaturation (95 C, 15 seconds, optics off) and annealing/extension (66 C, 60 seconds, optics on). The results of the modified realtime PCR were interpreted as follows: ‘detected’ if the cycle threshold score (CT) was 36 or lower, ‘indeterminate’ for CTs between 36 and 40, and ‘not detected’ for a CT of 40.

### Feeding trial

Two experimental groups (Exposed and Unexposed) of 20 lambs each were selected from the segregated ewe groups based on similar weaning weights and twin/single lambing types (Table 1), moved to Washington State University and housed in separate barns 90 m apart and that housed no other sheep or goats. Lambs from each group were allotted to lightweight and heavyweight pens of 10 lambs each based on initial weights. Pens were 62 to 78 m^2^ in area. The lambs were fed a pelleted alfalfa-barley ration for the trial, which was initially supplemented by long stemmed hay offered ad libitum during a one week transition period. The pellet composition (per 1000 kg) was alfalfa meal 870 kg), barley (100 kg), yellow grease (20 kg), ammonium chloride (5 kg), selenium-supplemented trace mineral salt (3.5 kg), lasalocid (1.8 kg at 12.43 g/kg; Bovatec, Zoetis, Parsippany NJ), vitamin A premix (0.04 kg at 30 KIU/g), vitamin D premix (0.03 kg at 8.8 MIU/kg), vitamin E-50 (0.02 kg at 500 KIU/kg), and Manganese Oxide (0.02 kg). The pelleted ration was determined to contain 16.9% crude protein, 36.9% neutral detergent fiber, 26.2% acid detergent fiber, 4.8% lignin/cutin, and 1.4% acid insoluble ash/silica (Dumas combustion method, AOAC 990.03, WSU Wildlife Habitat/Nutrition Laboratory, Pullman WA). Lambs remained on feed until a final target weight of 60kg was attained. Lambs were harvested in 3 groups at the WSU Meats Laboratory as they attained the target weight.

**Table 1:**
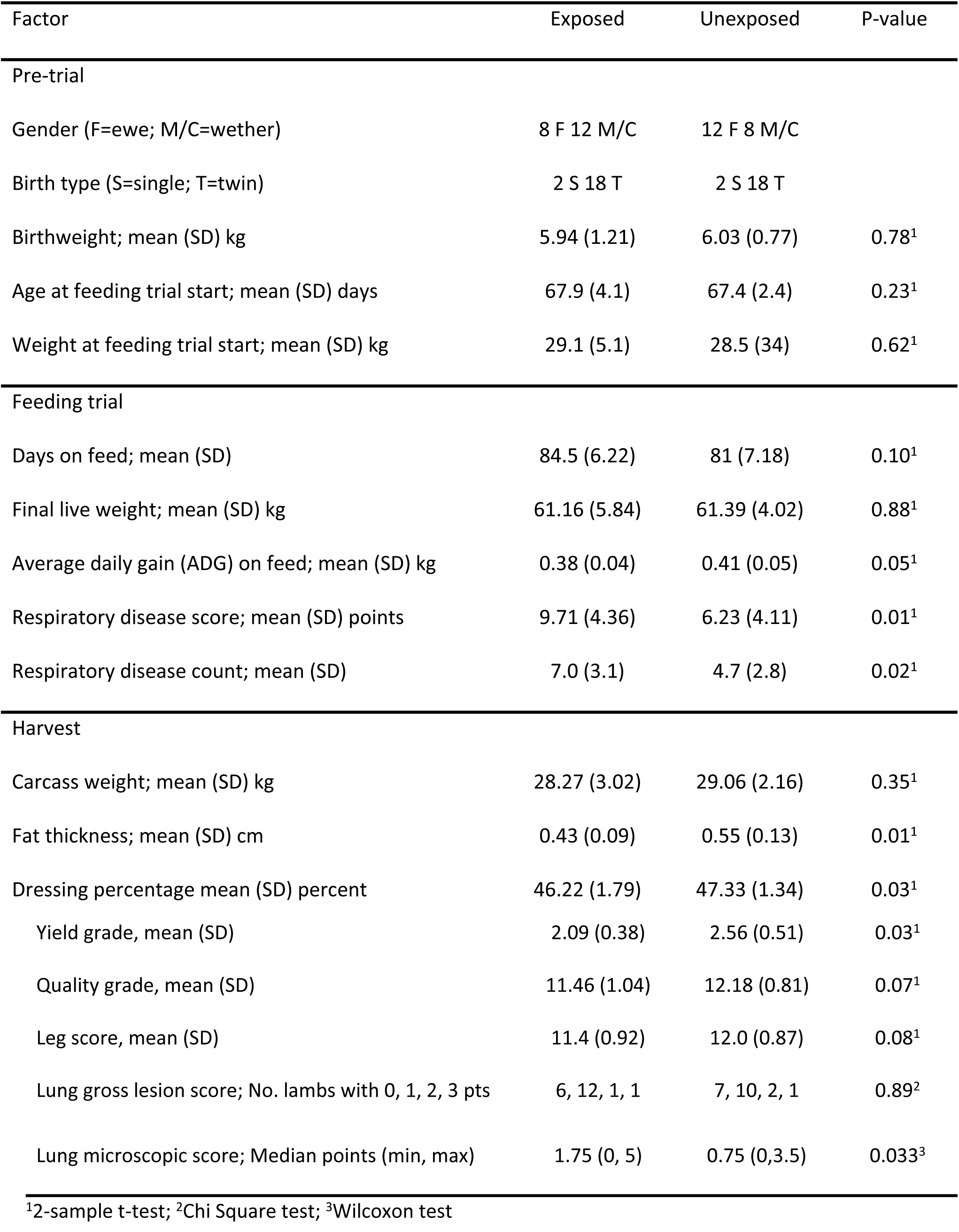
Demographic, growth, and carcass factors of lambs by *M. ovipneumoniae* exposure group prior to and during the feeding trial and at harvest.

### Health monitoring

Each lamb group was observed by a single scorer for 15 minute periods on each of 21 days between 28 and 74 days on feed to detect coughing, increased nasal discharge, nose licking, head shaking or increased respiratory rates. For each group on each observation day, the numbers of lambs observed with one or more of these signs, and the total number of signs observed were tallied. Because of the segregated nature of the barn containing the group of lambs with no exposure to *M. ovipneumoniae*, it was not possible to blind the observer to the treatment group identity.

### Carcass measurements

Hot carcass weights were obtained at harvest. At approximately 48 h postmortem, leg conformation score, adjusted 12th rib fat thickness, quality grade and yield grade based on USDA standards [21] were determined by an experienced lamb carcass evaluator (JB).

### Pathology

After plucks were removed from the lamb carcasses, the hearts were separated and the lungs and tracheas were removed from the kill floor, photographed dorsally and ventrally, and tissue samples from both anterior lung lobes were removed and fixed in 10% neutral buffered formalin. Gross lung photos were evaluated by a pathologist blind to the treatment group of the lambs, and scored on a scale of 0 to 3 representing observed consolidation affecting 0, <5%, 5-10%, and >10%, respectively, of the lung surface. After fixation, lung tissues were processed and embedded in paraffin, sectioned, and stained by hematoxyln and eosin using routine procedures at the Washington Animal Disease Diagnostic Laboratory. Slides were evaluated by a ACVP-certified veterinary pathologist (KAP) unaware of the exposure group source of the tissues.

## Results and Discussion

Prior to lambing in 2016, a group of commingled ewes was serially sampled to detect *M. ovipneumoniae* colonization using nasal swabs tested by a realtime PCR assay that targets a species-specific region of the small ribosomal subunit encoding locus. Heterogeneity of *M. ovipneumoniae* detection was observed, including some ewes that tested positive on every sampling date (carrier ewes) while others consistently tested negative. Our working definition of non-carrier ewes included those whose median CT score for the realtime PCR was 40, interpreted as ‘not detected’ (Figure 1).

**Figure 1:**
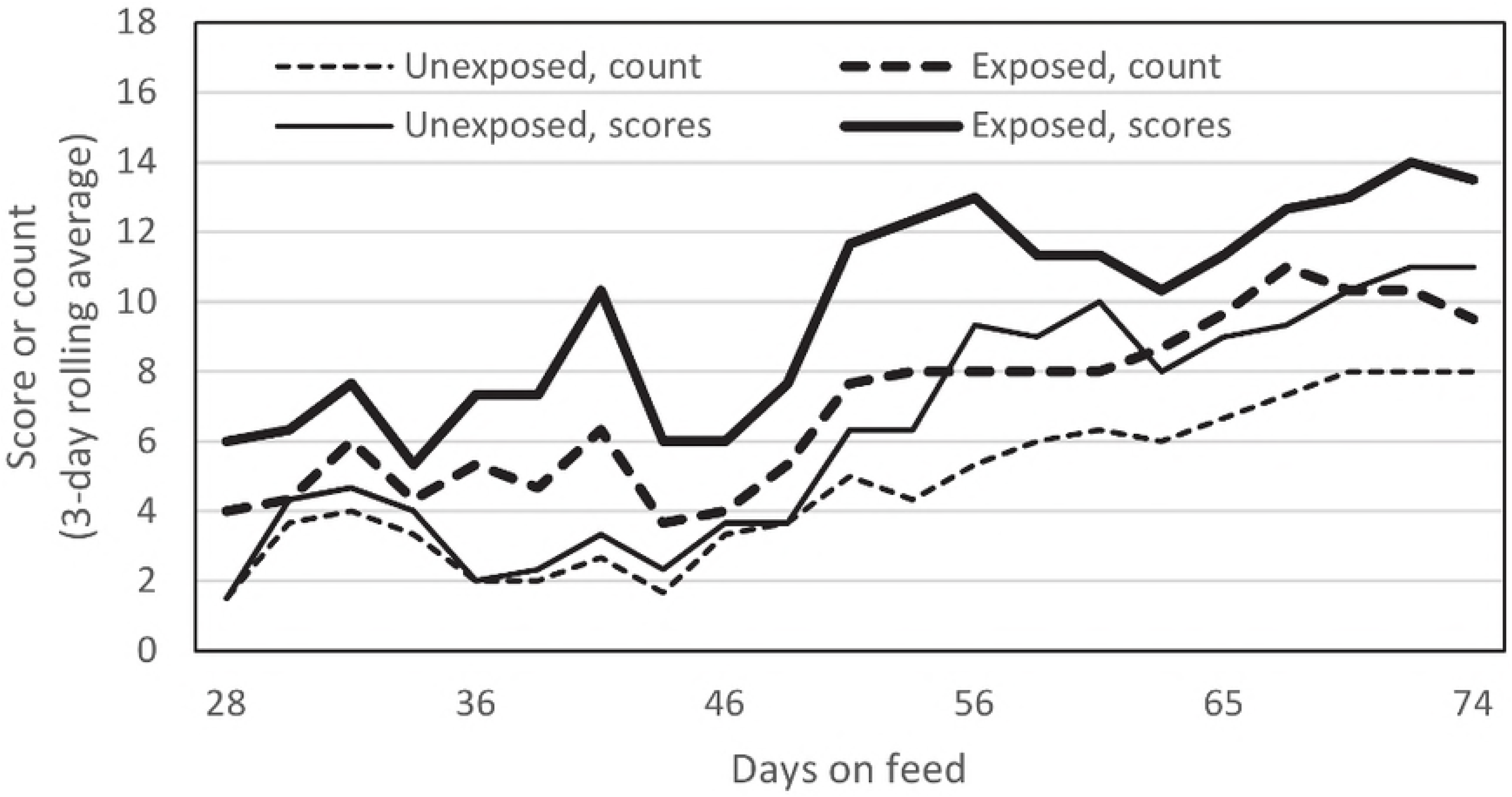
Results of longitudinal sampling of 34 ewes for *M. ovipneumoniae*, using nasal swabs tested by reatlime PCR. Each ewes was sampled on 5 or 6 occasions between November 2015 and March 2016. In each boxplot, the central heavy line represents the median value, the boxes represent the quartiles, the whiskers indicate the largest and smallest observations within 1.5 quartiles of the box, and circles represent outlying values. The realtime PCR test is interpreted based on the CT score, with scores below 36 interpreted as detection of *M. ovipneumoniae*, scores of 40 interpreted as non-detection, and scores between 36 and 40 interpreted as indeterminate for detection.

After lambing, non-carrier and carrier ewes, together with their lambs, were placed in separate pens that did not permit nose-to-nose contact. The non-carrier ewe group included 16 ewes with 27 lambs and the carrier ewe group included 18 ewes with 30 lambs. When the lambs reached approximately 60 days of age, *M. ovipneumoniae* was detected in a subset (7 of 21) of lambs in the carrier ewe pen, indicating the onset of natural vertical transmission of the agent; all non-carrier group lambs tested negative for *M. ovipneumoniae* at that time. At weaning at approximately 65 days of age, experimental groups of 20 lambs each were selected from the carrier and non-carrier pens, balancing lamb body weights and twin/single lambing types across the groups (Table 1), and moved to Washington State University, where they were housed in separate facilities separated by 90 m and transitioned to the pelleted ration over a one week period.

All lambs on trial were weighed and tested for *M. ovipneumoniae* carriage at 27, 46, 61, 74, and 88 days on feed. *M. ovipneumoniae* was never detected in any unexposed lamb during the feeding period, but was detected from all exposed lambs on each sampling date after the start of the trial. With regular observation, diverse respiratory disease signs were observed in both groups of lambs. However, signs were observed in more lambs per observation date, and more signs were observed per lamb in the exposed group than in the unexposed group (Table 1, Figure 2).

**Figure 2:**
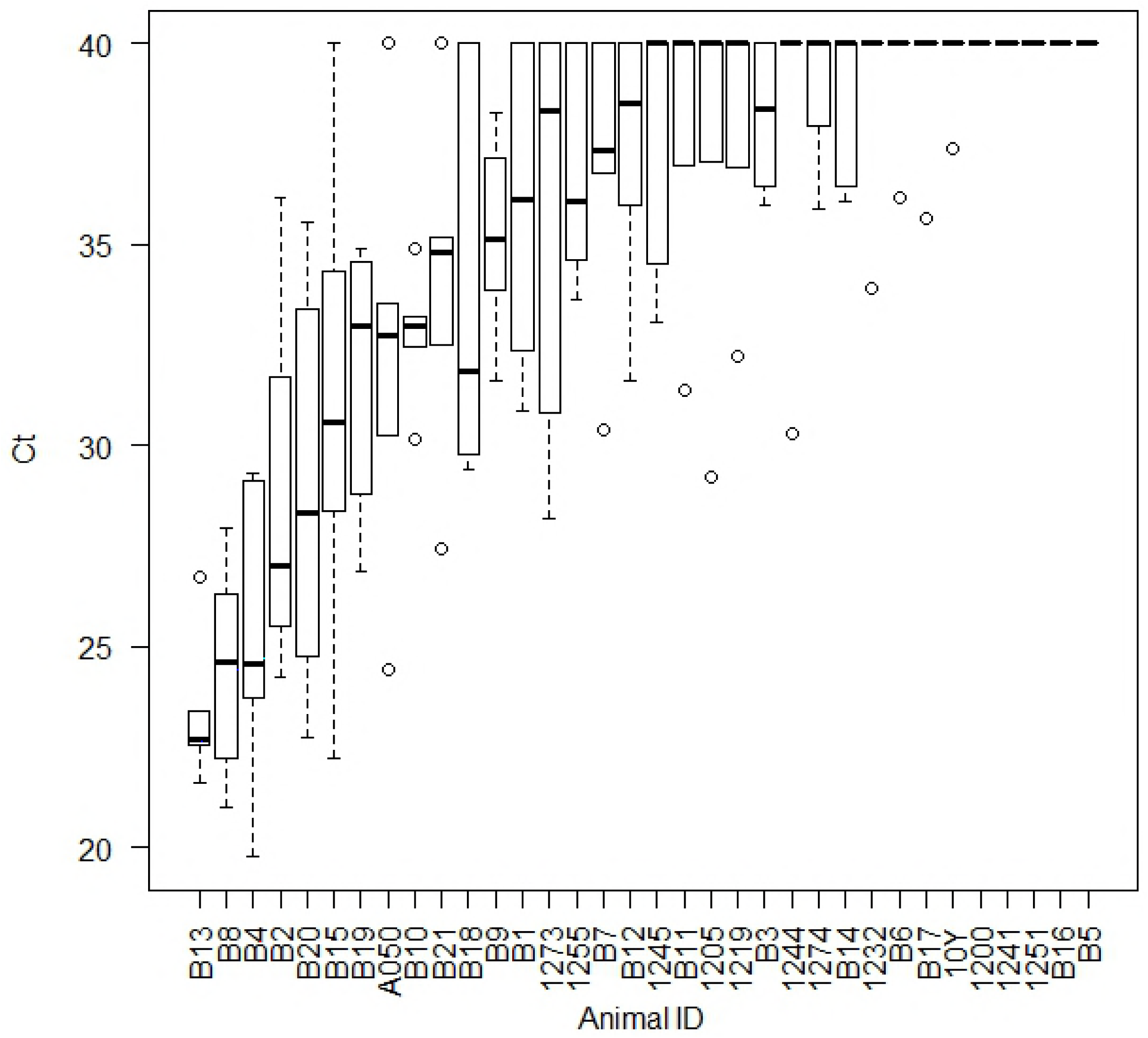
Cumulative counts of lambs exhibiting respiratory disease signs, and their cumulative respiratory signs scores. Data plotted represent 3-date rolling averages.

As they reached the target body weight, lambs were harvested in three groups at the WSU Meats Laboratory, including 15 lambs each after 74 and 88 days on feed. The final ten lambs, harvested on trial day 91, included several lambs that failed to reach the target weight. While most lambs reached 55 kg or higher body weight, two lambs in each group weighed <55 kg (Exposed group: 47.2, 45.9 kg; non-exposed group: 51.8 and 53.6 kg). At harvest, *M. ovipneumoniae* was not detected from bronchi of any unexposed group lambs. *M. ovipneumoniae* was detected in one or more bronchial swabs of all 20 exposed group lambs, including one bronchus of two lambs, two bronchi of one lamb, and all three bronchi of the remaining 17 lambs.

Univariate comparisons of lamb growth during the pre-feeding and -feeding periods, carcass traits, and gross and microscopic lung lesions are presented in Table 1. Factors that differed significantly between exposed and unexposed group lambs included average daily gains, respiratory disease scores and counts, carcass traits including fat thickness, dressing percentage, and yield grade, and microscopic lung lesion scores. Interestingly, the most pronounced group-associated differences in average daily gains occurred during the first month on feed, when average daily gains were 0.26 ± 0.16 (mean ± SD) for exposed lambs and 0.039 ± 0.13 for unexposed lambs (*P*=0.008, 2-sample t-test), coinciding with the period when the prevalence of *M. ovipneumoniae* infection increased from 33% to 100% of the exposed lamb group. Others have also observed that the most dramatic effects of experimentally reproduced chronic non-progressive pneumonia occur during the first 30 days after challenge [5, 6]. Analysis of variance was conducted using the model ADG ~ Group + Sex + Birth type + error. No significant 2-way interactions were detected. Independent significant effects of the *M. ovipneumoniae* exposure group and the birth number (single vs twin) on average daily gain (ADG) were detected, along with a trend for significance of lamb gender (Table 2).

**Table 2:** Analysis of variance of average daily gains (ADG) in kilograms through the duration of the feeding trial. Residual standard error: 0.043 on 36 degrees of freedom; Multiple R-squared: 0.2627, Adjusted R-squared: 0.2013; F-statistic: 4.275 on 3 and 36 DF, p-value: 0.011

| Factor* | ADG to harvest |  |  |  |
| --- | --- | --- | --- | --- |
|  | Estimate | Std. Error | t value | P-value |
| Intercept | 0.452 | 0.023 | 19.39 | <0.001 |
| Group (Exposed) | -0.034 | 0.014 | -2.42 | 0.021 |
| Sex (wether) | 0.0198 | 0.014 | 1.42 | 0.164 |
| Birth type (twin) | -0.056 | 0.023 | -2.46 | 0.019 |

\*Group (Exposed) in comparison with the *M. ovipneumoniae*-free group; Sex (wether) in comparison with ewe lambs; Birth type (twin) in comparison single births.

Our results confirm the feasibility of producing a cohort of lambs free from *M. ovipneumoniae* exposure. In this study, unexposed lambs exhibited fewer respiratory disease signs, gained body weight faster, tended to have improved carcass quality traits and had fewer microscopic lung lesions at harvest than the conventional *M. ovipneumoniae* exposed lambs; however, due to the single source of animals and the small numbers of study animals, health and growth effects need additional assessments in future studies to determine their repeatability.

The design of this trial differed from previous studies of the health and productivity effects of chronic non-progressive pneumonia of lambs in several ways. First, our design was based on the hypothesis that *M. ovipneumoniae* is a necessary component of this multi-factorial, polymicrobial disease. While we did observe subtle increases in clinical signs of respiratory disease and microscopic evidence of lung pathology in the lambs in the exposed group in this study, gross lesions of moderate or severe pneumonia were rare and did not differ among groups. This is an important observation because many previous studies of the adverse health and economic effects of chronic non-progressive pneumonia relied wholly or in part on detection of grossly visible lung lesions as the outcome variable [7-9, 22], suggesting that their findings could have been confounded by effects of sub-clinical *M. ovipneumoniae* infections in their ‘non-pneumonic’ controls. Secondly, the study flock where our lambs originated was not experiencing higher than average or expected respiratory disease losses at the time of the study, whereas some previous studies had selected flocks based on higher than expected frequency of pneumonia mortality or higher than average prevalence of lung lesions at necropsy [8, 22]. Thirdly, our method of generating the study groups ensured that the groups differed in exposure to *M. ovipneumoniae* throughout the duration of the trial, not just at the end, whereas previous trials may not have documented *M. ovipneumoniae* infection at all or accounted for infections that may have been cleared by the time of harvest. Together, these differences may have provided this study with better sensitivity to detect an effect of *M. ovipneumoniae* on lamb growth and productivity compared to previous efforts.

This study has several limitations, including the relatively small lamb group sizes, and the sourcing of all lambs from a single premises with a moderate prevalence of *M. ovipneumoniae* colonized ewes. Therefore, the results may not predict results on other premises with different geographical locations, biosecurity practices, ewe breeds, or prevalence of *M. ovipneumoniae* colonization. For example, on a high prevalence operation, it may be difficult or impossible to identify sufficient noncarrier ewes to produce a non-exposed lamb cohort of a desired size.

In conclusion, this pilot study documented the feasibility of producing a cohort of *M. ovipneumoniae-unexposed* animals through longitudinal application of a sensitive nasal swab – realtime PCR method. We also observed improved growth and carcass quality in the *M. ovipneumoniae-unexposed* lamb group, although those results require replication to confirm their magnitude and consistency. In general, these results support the hypothesis that elimination of *M. ovipneumoniae* may be feasible and may be of real economic value, providing a rationale for additional, larger scale studies.

